# Attenuated Salmonella-Mediated Delivery of GSDMD Potentiates PD-1 Blockade Therapy against Melanoma

**DOI:** 10.64898/2026.09.02.748795

**Authors:** Xiaolong Jia, Yingjie Yuan, Yujing Huang, Kairun Li, Xuejing Xu, Xiyu Zhang, Yu Zhang, Tiesuo Zhao, Lei Wang, Huijie Jia

## Abstract

Immunotherapy has emerged as a core therapeutic strategy for melanoma. Programmed death protein 1 (PD-1) is a critical immune checkpoint molecule that restrains host anti-tumor immunity, and therapeutic agents blocking the PD-1 signaling pathway have been widely deployed in clinical practice. Nevertheless, single-agent PD-1 blockade fails to elicit robust clinical responses in the majority of patients. Therefore, there is an urgent unmet need to develop combinatorial regimens capable of augmenting the anti-tumor efficacy of PD-1 inhibition. Gasdermin D (GSDMD), a pore-forming effector protein that orchestrates pyroptosis, exerts inherent anti-tumor activities upon overexpression. However, whether GSDMD can synergize with PD-1 blockade to enhance therapeutic outcomes against melanoma remains poorly defined. To address this question, we established an attenuated Salmonella engineered strain for targeted delivery of GSDMD, and further investigated the anti-melanoma therapeutic efficacy of combining this engineered bacterium with anti-PD-1 antibody via immunofluorescence staining, flow cytometry and other analytical approaches. Our in vivo results demonstrated that combinatorial treatment markedly suppressed melanoma progression in tumor-bearing mice relative to monotherapy with either GSDMD-expressing bacteria or anti-PD-1 antibody alone. Mechanistically, co-treatment upregulated intratumoral expression of GSDMD and the pro-apoptotic protein BAX, while simultaneously downregulating PD-1 expression. In addition, the GSDMD/anti-PD-1 combination significantly elevated the proportions of CD4⁺ and CD8⁺ T lymphocytes in both peripheral blood and splenic tissues, and facilitated robust tumor infiltration by these two T cell subsets. Compared with phosphate-buffered saline (PBS) and scramble control groups, combinatorial therapy promoted tumor infiltration of M1-type tumor-associated macrophages (TAMs) and repolarized TAMs away from the immunosuppressive M2 phenotype. Consistently, serum levels of the pro-inflammatory cytokines TNF-α and IFN-γ were markedly elevated following combined intervention. Collectively, this study verifies that attenuated Salmonellacarrying GSDMD synergizes with anti-PD-1 antibody to elicit potent anti-tumor effects in melanoma-bearing mice by amplifying systemic and intratumoral anti-tumor immune responses, which provides a preclinical rationale for novel combinatorial therapeutic strategies against melanoma.

## 1. Introduction

The global incidence of melanoma has been steadily rising, and melanoma ranks among the most aggressive subtypes of cutaneous malignancies. Although melanoma is only the third most prevalent form of skin cancer, it constitutes the leading cause of skin cancer-related mortality worldwide. Epidemiological statistics indicate that melanoma accounts for merely 4% of all cutaneous cancer cases yet is responsible for approximately 80% of deaths attributed to skin malignancies, highlighting its severe threat to human health. Furthermore, the global incidence and disease burden of melanoma have increased substantially over the past several decades [1–3]. Collectively, these statistics underscore an urgent clinical demand for the development of effective therapeutic regimens against melanoma.

Over the past decade, immunotherapy has revolutionized the therapeutic landscape of advanced melanoma and represented a paradigm shift in the field of oncology [4]. The advent of immune checkpoint inhibitors (ICIs) has marked a landmark breakthrough; compared with conventional therapeutic modalities, ICIs substantially improve survival outcomes in melanoma patients [5]. The remarkable clinical success of ICIs against melanoma has not only drastically transformed patient prognosis but also laid a foundational framework for expanding immunotherapeutic applications across multiple other malignancies. Long-term survival data from pivotal clinical trials have validated that ICI treatment elicits unprecedented durable therapeutic responses, enabling sustained disease control lasting several years in a subset of patients [6,7].

Programmed death-1 (PD-1), encoded by the PDCD1 gene, belongs to the immunoglobulin superfamily. The cytoplasmic tail of PD-1 contains two conserved tyrosine-based motifs: the immunoreceptor tyrosine-based inhibitory motif (ITIM) and the immunoreceptor tyrosine-based switch motif (ITSM). Cumulative evidence has verified that the ITSM motif is an indispensable structural domain required for PD-1 to exert immunosuppressive effects on activated T cells [8–10]. As an inhibitory T cell surface receptor, PD-1 specifically recognizes immunosuppressive signals triggered by programmed death-ligand 1 (PD-L1) overexpressed on malignant tumor cells. This interaction suppresses T cell effector functions and ultimately facilitates tumor immune evasion [11,12].

PD-1 checkpoint blockade exhibits outstanding therapeutic efficacy for advanced melanoma, driving tumor regression and conferring long-term disease stabilization. Approximately half of treated patients derive clinical benefits from this regimen, in stark contrast to the less than 10% response rate observed with traditional therapies. Accordingly, PD-1 blockade has emerged as a highly promising novel therapeutic modality for advanced melanoma [6].

Nevertheless, despite the landmark advances brought by PD-1 inhibitor-based immune checkpoint blockade, which have profoundly reshaped clinical outcomes for a subset of melanoma patients [13,14], prominent therapeutic limitations persist in real-world clinical practice. The majority of melanoma patients exhibit low clinical response rates to monotherapy with single-agent PD-1 inhibitors, and a proportion of patients fail to derive any therapeutic benefit at all. Accordingly, developing rational combinatorial regimens that harness synergistic effects between distinct therapeutic modalities to overcome the drawbacks of PD-1 monotherapy has emerged as a pivotal research priority and core strategy to amplify the efficacy of PD-1 blockade and improve overall prognosis in melanoma patients [15,16].

Pyroptosis, an inflammatory form of programmed cell death, has recently been established as a critical cellular biological process implicated in numerous diseases including malignancies. Accumulating preclinical evidence has validated that targeting pyroptotic signaling cascades augments the anti-tumor potency of conventional cancer therapies and reinforces cytotoxicity against drug-resistant tumor cells, offering novel therapeutic avenues to tackle treatment resistance in malignant tumors [17,18]. Emerging studies have uncovered a functional crosstalk between pyroptosis and anti-tumor immunity, implying that pyroptosis induction can trigger robust systemic anti-tumor immune responses and serve as a promising adjuvant strategy to optimize cancer treatment. To date, an expanding body of research has investigated the synergistic potential of combining pyroptosis-inducing interventions with immune checkpoint blockade for cancer management, which lays a solid translational foundation for combinatorial tumor immunotherapy [19,20].

Gasdermin D (GSDMD), the core executioner protein of pyroptosis, undergoes proteolytic cleavage by inflammasome-activated caspase-1 as well as lipopolysaccharide (LPS)-triggered caspase-11/4/5. Cleaved GSDMD subsequently participates in the pathological progression of a wide spectrum of disorders and has emerged as a promising therapeutic target for cancer and other diseases [21]. Previous research has demonstrated that cucurbitacin B, a bioactive phytochemical derived from natural plants, selectively acts on non-small cell lung cancer (NSCLC) cells by triggering GSDMD-dependent pyroptosis. This cascade robustly suppresses cancer cell proliferation and viability, exerts potent anti-lung tumor activity, and ultimately halts NSCLC progression [22].

In the context of melanoma, CRKL—a vital member of the chicken tumor virus number 10 oncogene homolog (CRK) protein family—functions as a central regulator of cellular signal transduction and oncogenesis. Knockdown of CRKL elevates endogenous GSDMD expression, thereby repressing melanoma cell proliferation and invasive capacity and sensitizing melanoma cells to cisplatin-based chemotherapy [23]. However, it remains largely unelucidated whether enforced GSDMD expression can potentiate the anti-melanoma efficacy of PD-1 blockade.

To address this unmet question, the present study investigated the anti-melanoma efficacy of combining an attenuated Salmonella strain—an original genetically engineered vector constructed in our laboratory to deliver the GSDMD expression cassette—with anti-PD-1 antibody. Our findings provide preclinical evidence and novel research insights for developing more potent combinatorial regimens that augment the therapeutic performance of PD-1 blockade against melanoma.

## 2. Materials and Methods

### 2.1 Animal Grouping and Therapeutic Regimen

Subcutaneous melanoma-bearing mouse models were established via inoculation of B16 melanoma cells. On day 7 post tumor inoculation, all mice were randomly assigned into five experimental groups: PBS group, Scramble group, anti-PD-1 monotherapy group, GSDMD monotherapy group, and anti-PD-1 + GSDMD combination group. The treatment regimens for each group were specified as follows:

PBS group: intratumoral injection of 100 μL phosphate-buffered saline (PBS); Scramble group: intratumoral administration of 4 × 10⁵ CFU attenuated engineered Salmonella carrying scramble sequences;

Anti-PD-1 group: intraperitoneal injection of 200 μg anti-PD-1 antibody;

GSDMD group: intratumoral delivery of 4 × 10⁵ CFU attenuated engineered Salmonella harboring the GSDMD expression plasmid;

Combination group: co-treatment with intraperitoneal 200 μg anti-PD-1 antibody plus intratumoral 4 × 10⁵ CFU attenuated engineered Salmonella carrying the GSDMD expression plasmid.

Anti-PD-1 antibody was administered once every two days for a total of four doses. Attenuated engineered Salmonella strains were injected once every seven days, with two doses in total. All mice were humanely sacrificed five days after the final treatment, followed by collection and detection of indicators including tumor weight.

### 2.2 One-step TUNEL Assay for Evaluating the Combined Effects of GSDMD-Expressing Attenuated Salmonella and Anti-PD-1 Antibody on Tumor Tissue Apoptosis

Prepared tumor tissue sections were subjected to deparaffinization and rehydration firstly. Subsequently, DNase-free proteinase K working solution (20 μg/mL diluted in 10 mM Tris-HCl buffer, pH = 7.4) was added onto tissue sections, followed by incubation at 37 °C for 60 min. After incubation, sections were rinsed three times with phosphate-buffered saline (PBS), with 5 min for each wash. According to the manufacturer’s instructions of the TUNEL detection kit, sections were covered with equilibration buffer and incubated at room temperature (RT) for 10 min. After discarding the equilibration buffer, TUNEL reaction mixture was supplemented to fully cover the tissue specimens. All sections were placed in a humidified chamber and incubated at 37 °C for 60 min under dark conditions to avoid fluorescence quenching. The reaction was terminated by three rounds of PBS rinsing, and each washing step lasted for 5 min. Thereafter, sections were counterstained with DAPI staining solution for 10 min at RT. After adequate PBS washing, slides were mounted with anti-fade mounting medium. Ultimately, fluorescence observation and image acquisition were performed under a laser scanning confocal microscope. The FITC green fluorescent signal was detected at an excitation wavelength of 488 nm, and apoptotic nuclei were characterized by positive green fluorescence.

### 2.3 Western Blot Analysis of Tumor-Associated Protein Expression Modulated by GSDMD-Expressing Attenuated Salmonella Combined with Anti-PD-1 Antibody

Tumor tissues were dissected from melanoma-bearing mice and immediately preserved in an ultra-low temperature refrigerator at −80 °C for subsequent experiments. For protein extraction, 0.1 g of tumor tissues from each group were weighed precisely and homogenized thoroughly using a tissue grinder with pre-cooled protein lysis buffer. After centrifugation, the supernatant was collected as total tumor protein extracts. The expression levels of target proteins were quantified via Western blot assay, and the detailed experimental procedures were described as follows:

SDS-PAGE resolving gels and stacking gels with optimized concentrations were prepared, and equal amounts of protein samples were loaded for electrophoresis to separate proteins with distinct molecular weights. After electrophoresis, separated protein bands were electrotransferred onto PVDF membranes. Subsequently, 5% skimmed milk blocking solution was prepared, and membranes were incubated in the blocking solution at room temperature for 2 h.

During the blocking period, primary antibodies against GSDMD, PD-1 and Cleaved-caspase-1 were diluted in accordance with the manufacturer’s protocols. Upon the completion of blocking, PVDF membranes were trimmed based on the prestained protein marker bands. After thorough rinsing with washing buffer, the trimmed membranes were incubated with corresponding diluted primary antibodies overnight at 4 °C. On the next day, membranes were washed sufficiently, followed by incubation with species-matched secondary antibodies at room temperature for 1 h. After another round of rinsing, ECL chemiluminescence substrate was evenly covered on membrane surfaces. Target protein bands were visualized and detected using a chemiluminescence imaging system.

### 2.4 Immunofluorescence Assay for Detecting Protein Expression and Immune Cell Infiltration in Tumor Tissues of Melanoma-Bearing Mice

Prepared tumor tissue sections were sequentially subjected to baking, deparaffinization and gradient rehydration, followed by heat-induced antigen retrieval. Briefly, sections were incubated in hot antigen retrieval buffer for 15 min per run, and the retrieval procedure was repeated twice. All sections were cooled naturally to room temperature (RT) afterwards. Subsequently, the sections were rinsed with PBS, and redundant liquid was wiped off carefully. Peroxidase blocking solution was added onto sections, followed by 10 min of incubation at RT to block endogenous peroxidase activity.

After three rounds of PBS washing, tissue sections were blocked with antigen blocking buffer for 10 min at RT. During the blocking process, primary antibodies against Ki-67, CD3, CD4, CD8, CD11b, CD86, CD206 and Granzyme B were diluted in strict accordance with the manufacturer’s instructions. Upon blocking completion, the blocking buffer was discarded, and diluted primary antibodies were applied to tissue sections. The slides were placed in a light-proof humidified chamber and incubated overnight at 4 °C.

On the next day, sections were equilibrated at RT for no less than 1 h. After sufficient PBS rinsing, species-matched secondary antibodies diluted per official protocols were added, and sections were incubated for 30 min at RT under light-shielded conditions. After another washing step, sections were counterstained with DAPI solution for 5 min in the dark. Finally, the slides were rinsed thoroughly with PBS, dehydrated slightly and mounted with anti-fade mounting medium. All samples were coverslipped, and fluorescence images were captured and visualized via a laser scanning confocal microscope.

### 2.5 Flow Cytometry Analysis of Immune Cell Proportions in Peripheral Blood and Spleen of Tumor-Bearing Mice Treated with GSDMD-Expressing Attenuated Salmonella Combined with Anti-PD-1 Antibody

Following orbital blood collection, mice were euthanized via cervical dislocation. Spleens were isolated aseptically and placed in petri dishes supplemented with RPMI 1640 medium. Fresh spleens were ground gently with frosted glass slides to release immune cells sufficiently, and the cell suspension was rinsed and collected with RPMI 1640 medium. Subsequently, the cell suspension was filtered through a 200-mesh cell sieve into 15 mL centrifuge tubes and placed on ice temporarily.

After sample volume balancing, cell suspensions were centrifuged at 1200 rpm for 5 min at 4 °C, and the supernatants were discarded to harvest cell pellets. Both splenocyte and peripheral blood cell pellets were resuspended in 4 mL erythrocyte lysis buffer and incubated on ice for 8 min for erythrocyte cleavage. Lysis reaction was terminated by adding 8 mL RPMI 1640 medium, followed by centrifugation at 1200 rpm for 5 min at 4 °C. The erythrocyte lysis procedure was repeated if cell pellets remained visibly red. After removing the supernatant, cell pellets were resuspended in 1 mL culture medium for cell counting, and a total of 1×10⁶ cells per sample were harvested for flow cytometry staining. Cell suspensions were transferred into 1.5 mL centrifuge tubes and centrifuged at 1200 rpm for 5 min at 4 °C, then the supernatants were discarded. Meanwhile, blank control tubes and single-stained compensation tubes were prepared for flow cytometry calibration.

Mixed antibody working solution containing CD3, CD4, CD8 and NK cell antibodies was diluted with RPMI 1640 medium at a dilution ratio of 1:200. Cell pellets were resuspended with 50 μL prepared antibody mixture, while the blank control group received no antibodies and compensation tubes were incubated with a single fluorescent antibody separately. All samples were incubated for 30 min at 4 °C under light-shielded conditions. Post incubation, each tube was supplemented with 500 μL RPMI 1640 medium, centrifuged and decanted. Cells were washed again with 300 μL medium, centrifuged and prepared for subsequent on-machine detection.

### 2.6 Enzyme-Linked Immunosorbent Assay (ELISA) Detection of Circulating Cytokine Levels in Tumor-Bearing Mice

The serum concentrations of TNF-α and IFN-γ were quantified using a double-antibody sandwich ELISA kit in strict accordance with the manufacturer’s instructions. Briefly, serially diluted standard reagents and serum samples from each experimental group were added into antibody-precoated microplates for incubation. After washing steps, biotinylated detection antibodies were supplemented, followed by another round of incubation and plate washing. Subsequently, horseradish peroxidase (HRP)-conjugated avidin was added to each well. After chromogenic reaction under light-shielded conditions, stop buffer was pipetted to terminate the reaction. The optical density (OD) value was measured at an excitation wavelength of 450 nm, and the final cytokine concentrations of serum samples were calculated based on the fitted standard curve.

### 2.7 Statistical Analysis

All experimental data were statistically analyzed using SPSS version 22.0 software. Intergroup differences were compared via one-way analysis of variance (ANOVA) and two-tailed unpaired Student’s t-test, respectively. All quantitative results were presented as the mean ± standard deviation (SD). A P-value less than 0.05 was defined as statistically significant.

## 3. Results

### 3.1 Combined Treatment with GSDMD-Expressing Attenuated Salmonella and Anti-PD-1 Antibody Efficiently Suppresses Melanoma Growth

To clarify the anti-tumor efficacy of GSDMD-engineered Salmonella combined with PD-1 blockade, melanoma-bearing mouse models were established, and tumor growth as well as tumor weight were monitored and analyzed (Figure 1A). Tumor morphological observation and statistical quantification demonstrated that compared with the PBS and Scramble groups, single-agent treatment with either GSDMD-expressing attenuated Salmonella or anti-PD-1 antibody remarkably retarded melanoma proliferation and reduced tumor weight. Notably, combination therapy exerted the most potent tumor-suppressive effect, triggering the lowest tumor weight among all experimental groups (Figure 1B–1C). Collectively, these findings indicated that the combinatorial regimen of GSDMD-delivering attenuated Salmonella plus PD-1 blockade serves as a promising therapeutic strategy against melanoma.

**Fig. 1.**
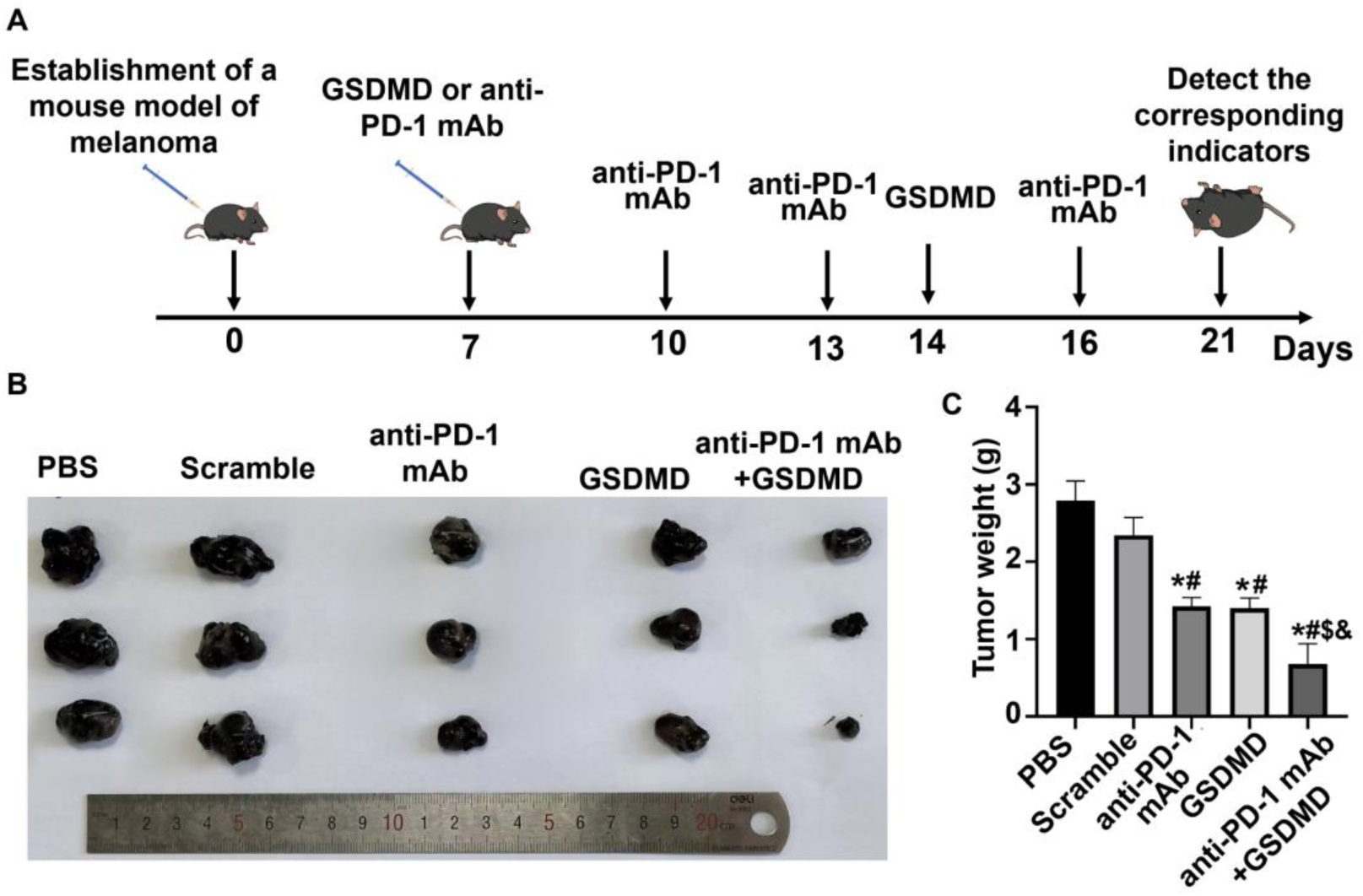
Effects of combined treatment with GSDMD-expressing attenuated Salmonella and anti-PD-1 antibody on melanoma growth (A) Schematic diagram illustrating the treatment schedule for GSDMD-expressing attenuated Salmonella plus anti-PD-1 antibody intervention. (B–C) Representative photographs of melanoma xenografts and statistical quantification of tumor weight acquired at day 5 after the final treatment. \**P* < 0.05 versus PBS group; ^#^*P* < 0.05 versus Scramble group; ^$^*P* < 0.05 versus anti-PD-1 monotherapy group; ^&^*P* < 0.05 versus GSDMD monotherapy group.

### 3.2 Combination Therapy Inhibits Tumor Cell Proliferation and Facilitates Tumor Cell Apoptosis in Melanoma Tissues

To further explore the regulatory effects of combined intervention on tumor proliferation, immunofluorescence staining was performed to detect the expression of Ki-67, a canonical proliferation biomarker. The results revealed that the expression level of intratumoral Ki-67 was markedly downregulated in both GSDMD monotherapy and anti-PD-1 monotherapy groups relative to the PBS and Scramble control groups. Moreover, combined treatment produced a superior inhibitory effect on Ki-67 expression (Figure 2A, 2C).

**Fig. 2.**
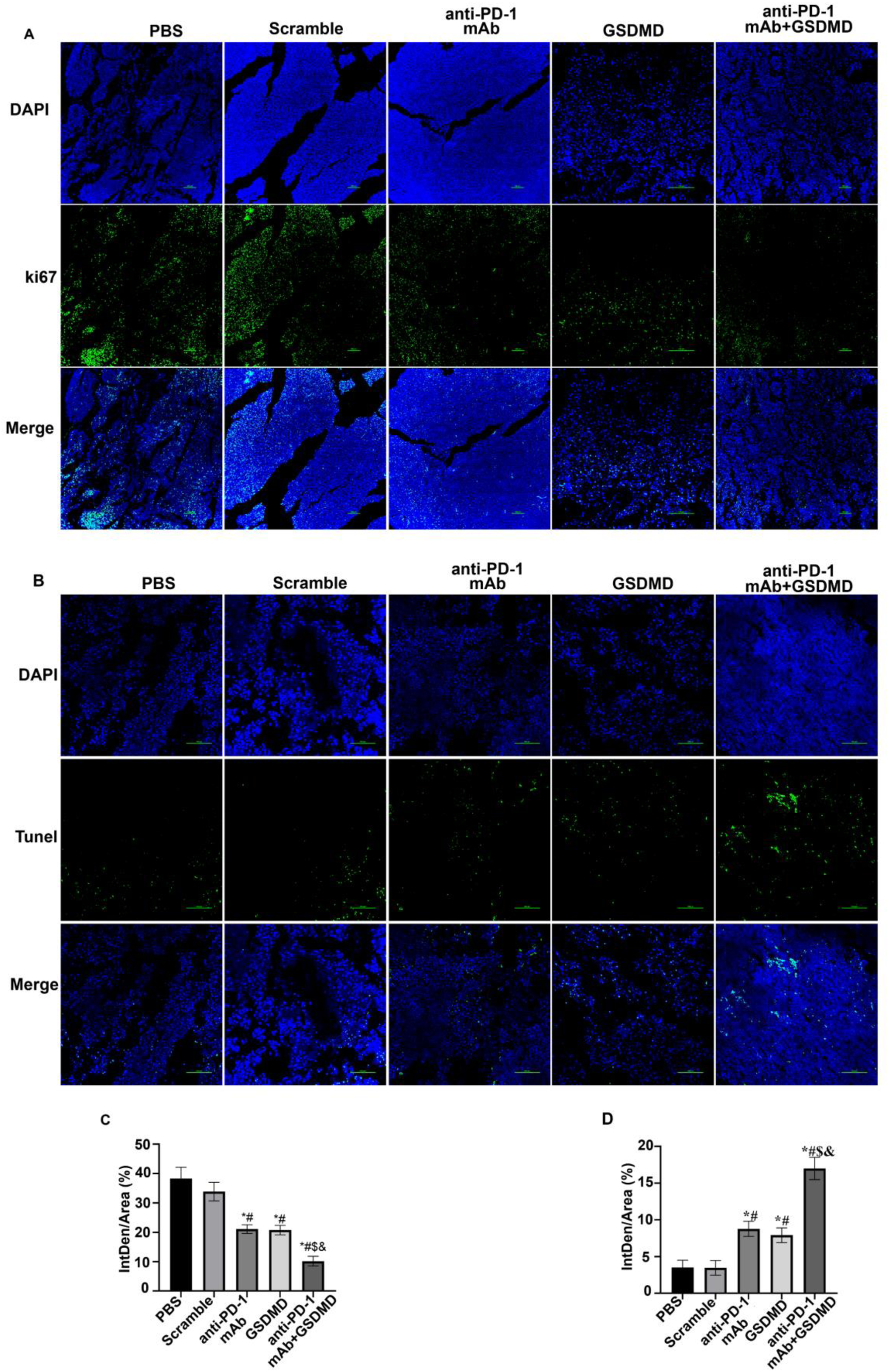
Combined GSDMD-expressing attenuated Salmonella and PD-1 blockade regulates cell proliferation and apoptosis in melanoma tissues (A, C) Immunofluorescence staining and corresponding quantitative statistics of intratumoral Ki-67 expression following combined treatment. (B, D) One-step TUNEL assay results and statistical analysis of apoptotic cell rates in tumor tissues upon co-administration of GSDMD-expressing attenuated Salmonella and anti-PD-1 antibody. \**P* < 0.05 versus PBS group; ^#^*P* < 0.05 versus Scramble group; ^$^*P* < 0.05 versus anti-PD-1 monotherapy group; ^&^*P* < 0.05 versus GSDMD monotherapy group.

In addition, the one-step TUNEL assay was conducted to evaluate tumor cell apoptosis across different groups. Consistent with the proliferation-related results, monotherapy with either GSDMD-expressing Salmonella or anti-PD-1 antibody moderately elevated the apoptotic rate of intratumoral cells compared with two control groups. Of note, co-administration of the two therapies further amplified tumor cell apoptosis (Figure 2B, 2D).

### 3.3 Effects of Combined Therapy with GSDMD-Expressing Attenuated Salmonella and PD-1 Blockade on Intratumoral Protein Expression

Subsequently, western blot assay was performed to quantify the intratumoral expression levels of GSDMD and PD-1. As indicated by the results, compared with the PBS and Scramble control groups, monotherapy with either GSDMD-expressing attenuated Salmonella or anti-PD-1 antibody markedly upregulated tumoral GSDMD expression. In addition, mice receiving GSDMD-engineered Salmonella exhibited higher GSDMD abundance than those treated with anti-PD-1 monotherapy. Notably, the combination group achieved the maximal GSDMD expression level among all groups (Figure 3A–3B).

**Fig. 3.**
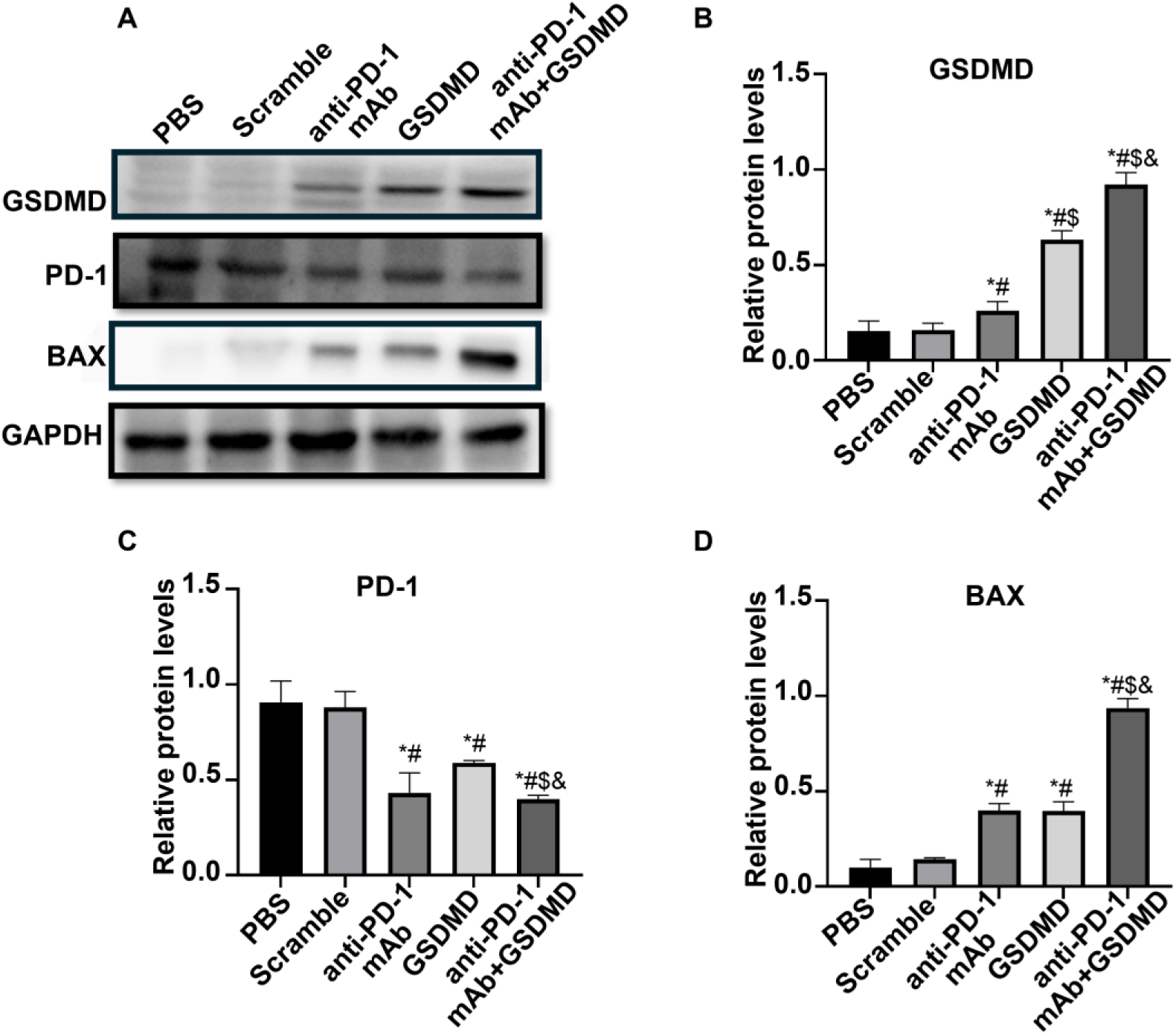
Impacts of combinatorial therapy with GSDMD-expressing attenuated Salmonella and anti-PD-1 antibody on intratumoral protein expression (A–D) Western blot detection and quantitative statistical analysis of GSDMD, PD-1 and BAX protein levels in melanoma tissues after PD-1 blockade combined with GSDMD-delivering attenuated Salmonella. \**P* < 0.05 versus PBS group; ^#^*P* < 0.05 versus Scramble group; ^$^*P* < 0.05 versus anti-PD-1 monotherapy group; ^&^*P* < 0.05 versus GSDMD monotherapy group.

For PD-1 expression profiling, anti-PD-1 monotherapy significantly inhibited intratumoral PD-1 expression relative to control groups. Surprisingly, GSDMD-expressing Salmonella alone also exerted an inhibitory effect on PD-1 expression. Furthermore, combined intervention triggered the lowest PD-1 protein abundance across all experimental groups (Figure 3A, 3C).

We further detected the expression of pro-apoptotic protein BAX to validate tumor apoptosis. Consistent with the apoptotic results obtained from one-step TUNEL assay, BAX expression was remarkably elevated in both GSDMD and anti-PD-1 monotherapy groups compared with the PBS and Scramble groups. More importantly, combination therapy induced a more pronounced upregulation of BAX expression than either single treatment (Figure 3A, 3D).

### 3.4 Combination of GSDMD-Expressing Attenuated Salmonella and PD-1 Blockade Increases Splenic T Lymphocyte Proportions in Tumor-Bearing Mice

Changes in splenic T lymphocyte subsets are essential indicators reflecting systemic anti-tumor immune function. Herein, flow cytometry was performed to quantify the proportion of T lymphocytes isolated from mouse spleens. The results revealed that both GSDMD-expressing attenuated Salmonella monotherapy and anti-PD-1 antibody monotherapy moderately elevated the percentages of splenic CD4⁺ T cells (Figure 4A, 4C) and CD8⁺ T cells (Figure 4B, 4D) relative to control groups. Of note, combined treatment further augmented the abundance of these two T-cell subsets.

**Fig. 4.**
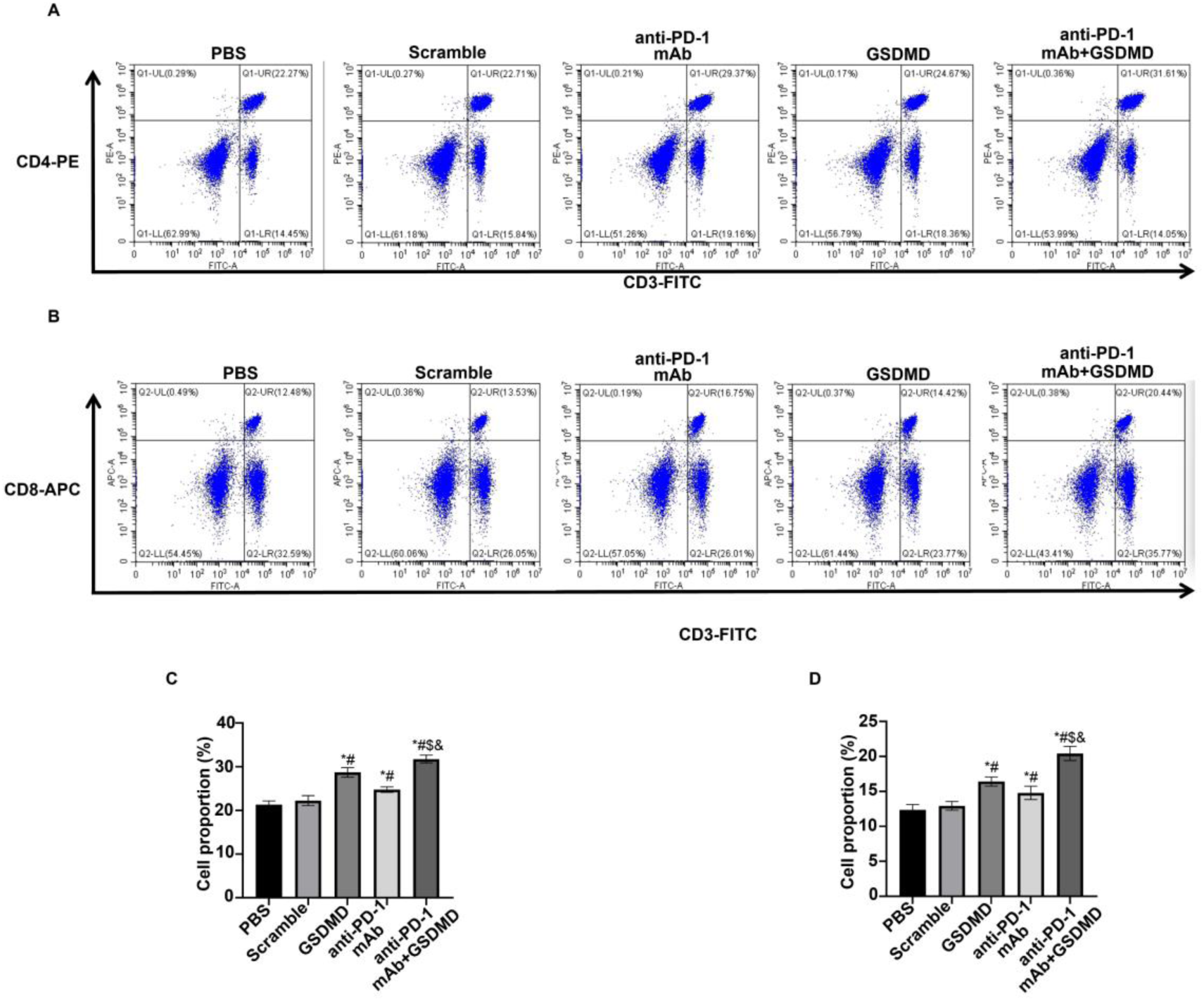
Effects of combined treatment with GSDMD-expressing attenuated Salmonella and PD-1 blockade on T lymphocyte proportions in mouse spleens (A, C) Flow cytometric detection and quantitative statistical analysis of splenic CD4⁺ T cell proportions after co-treatment with GSDMD-expressing attenuated Salmonella and anti-PD-1 antibody. (B, D) Flow cytometry and statistical quantification of splenic CD8⁺ T cell proportions following the combinatorial regimen. \**P* < 0.05 versus PBS group; ^#^*P* < 0.05 versus Scramble group; ^$^*P* < 0.05 versus anti-PD-1 monotherapy group; ^&^*P* < 0.05 versus GSDMD monotherapy group.

### 3.5 Combined Therapy Markedly Promotes Intratumoral Infiltration of T Lymphocytes and Granzyme B⁺ Cells

Immunofluorescence staining was conducted to evaluate the effects of co-treatment on tumoral CD4⁺ and CD8⁺ T-cell infiltration. Compared with the PBS and Scramble groups, monotherapy with either GSDMD-engineered Salmonella or anti-PD-1 antibody significantly raised intratumoral CD4⁺ T-cell infiltration, and the combination group exhibited the highest infiltration level of CD4⁺ T cells (Figure 5A–5B). Similarly, combined intervention exerted the most prominent effect on facilitating CD8⁺ T-cell accumulation within melanoma tissues (Figure 5C–5D).

**Fig. 5.**
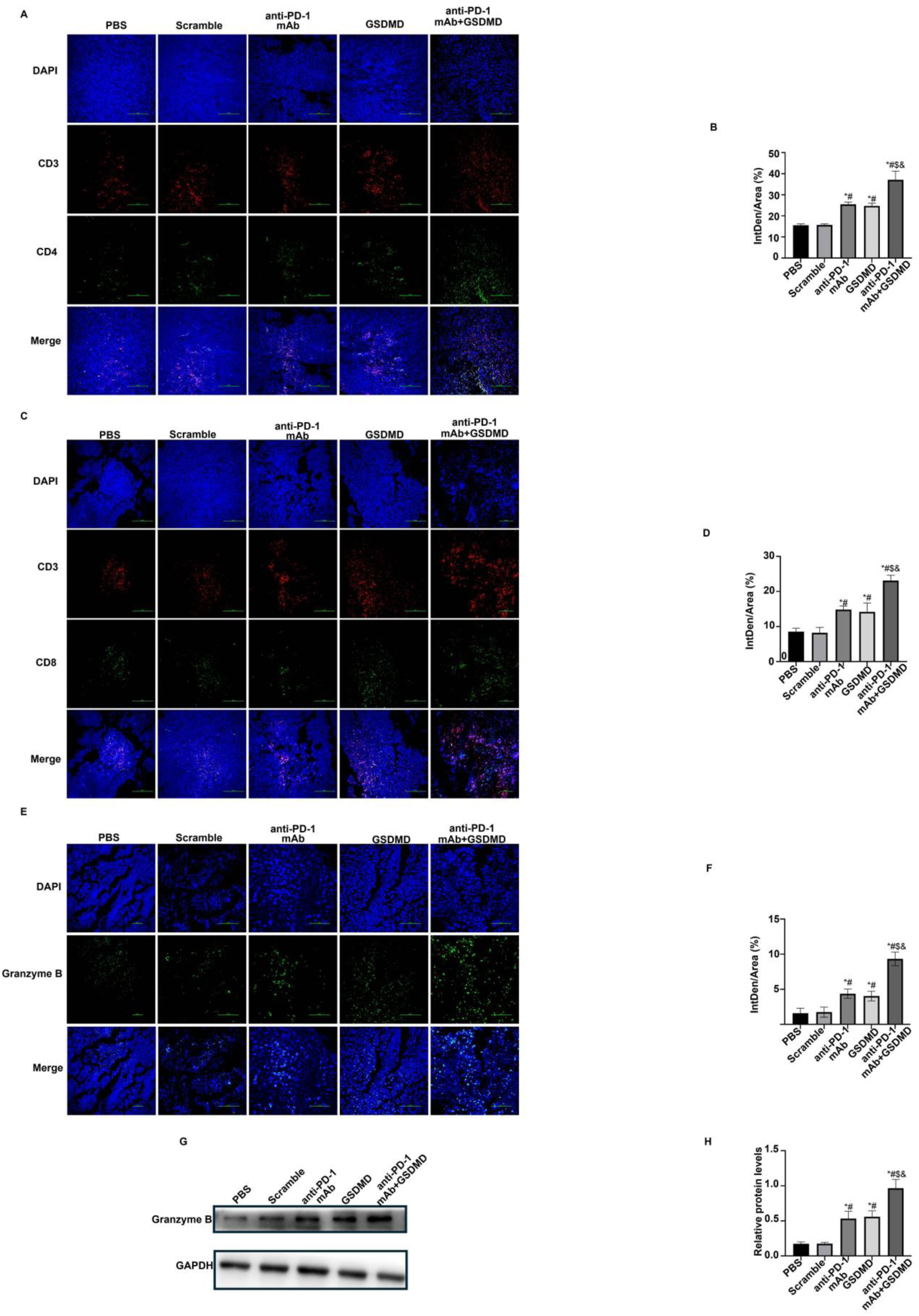
Combined GSDMD-expressing attenuated Salmonella and anti-PD-1 antibody enhances intratumoral infiltration of T lymphocytes and Granzyme B⁺ cells (A–B) Immunofluorescence staining and statistical analysis for the infiltration ratio of intratumoral CD4⁺ T cells under combinatorial intervention. (C–D) Immunofluorescence detection and quantitative statistics of intratumoral CD8⁺ T cell infiltration. (E–F) Immunofluorescence assay and corresponding quantification of intratumoral Granzyme B expression levels. (G–H) Western blot analysis and statistical quantification of Granzyme B protein abundance in melanoma tissues after combined therapy. \**P* < 0.05 versus PBS group; ^#^*P* < 0.05 versus Scramble group; ^$^*P* < 0.05 versus anti-PD-1 monotherapy group; ^&^*P* < 0.05 versus GSDMD monotherapy group.

Granzyme B is a core cytotoxic molecule that empowers effector T cells to eliminate malignant tumor cells. We further investigated whether this combinatorial regimen could modulate intratumoral Granzyme B expression. Immunofluorescence results demonstrated that either monotherapy efficiently upregulated Granzyme B expression compared with two control groups, while combinatorial treatment induced maximal Granzyme B accumulation in tumor tissues (Figure 5E–5F). Western blot assays further verified identical expression trends (Figure 5G–5H).

### 3.6 Combination of GSDMD-Expressing Attenuated Salmonella and PD-1 Blockade Facilitates Intratumoral M1 Macrophage Infiltration While Repressing M2 Macrophage Polarization

Tumor-associated macrophages exert dual regulatory effects on melanoma progression. Classically activated M1 macrophages elicit anti-tumor activities via tumor cell phagocytosis and antigen presentation, whereas alternatively activated M2 macrophages facilitate tumor proliferation, invasion and metastasis. CD86 serves as a canonical surface biomarker to reflect M1 macrophage activation, while CD206 is a specific marker of M2 macrophages. Accordingly, immunofluorescence staining was performed to assess the infiltration of CD86⁺ and CD206⁺ macrophages in tumor tissues. The results revealed that compared with the PBS and Scramble control groups, monotherapy with either GSDMD-expressing attenuated Salmonella or anti-PD-1 antibody increased intratumoral CD86⁺ macrophage infiltration (Figure 6A, 6C) and reduced the proportion of CD206⁺ macrophages (Figure 6B, 6D). Consistently, combined treatment further enhanced the accumulation of CD86⁺ macrophages and diminished CD206⁺ macrophage infiltration in melanoma tissues.

**Fig. 6.**
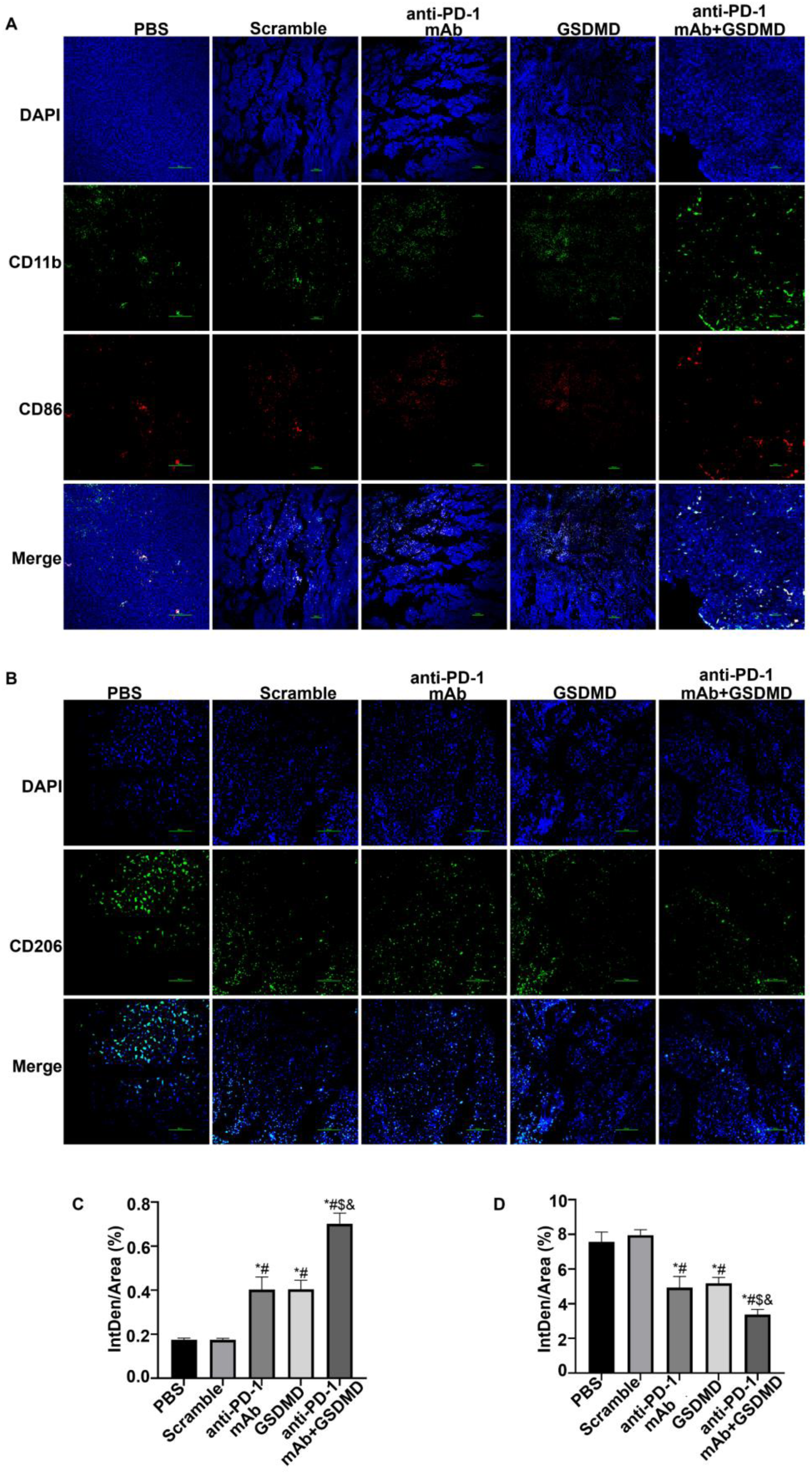
Impacts of GSDMD-expressing attenuated Salmonella combined with PD-1 blockade on the polarization ratio of tumor-associated macrophages (A, C) Immunofluorescence staining and statistical analysis of intratumoral CD86⁺ M1 macrophage infiltration. (B, D) Immunofluorescence detection and quantitative statistics of intratumoral CD206⁺ M2 macrophage infiltration. \**P* < 0.05 versus PBS group; ^#^*P* < 0.05 versus Scramble group; ^$^*P* < 0.05 versus anti-PD-1 monotherapy group; ^&^*P* < 0.05 versus GSDMD monotherapy group.

### 3.7 Combined Therapy Elevates Peripheral T Lymphocyte Proportions and Serum TNF-α/IFN-γ Levels in Tumor-Bearing Mice

Dynamic alterations in peripheral blood T lymphocyte subsets indirectly reflect systemic anti-tumor immune status, which is tightly correlated with tumor immune evasion and disease progression. Herein, flow cytometry was conducted to evaluate peripheral T lymphocyte proportions. Flow cytometric diagrams and statistical analysis demonstrated that the percentages of peripheral CD4⁺ T cells were markedly elevated in both GSDMD and anti-PD-1 monotherapy groups relative to control groups. Moreover, combinatorial treatment further upregulated CD4⁺ T cell abundance in peripheral blood (Figure 7A, 7C).

**Fig. 7.**
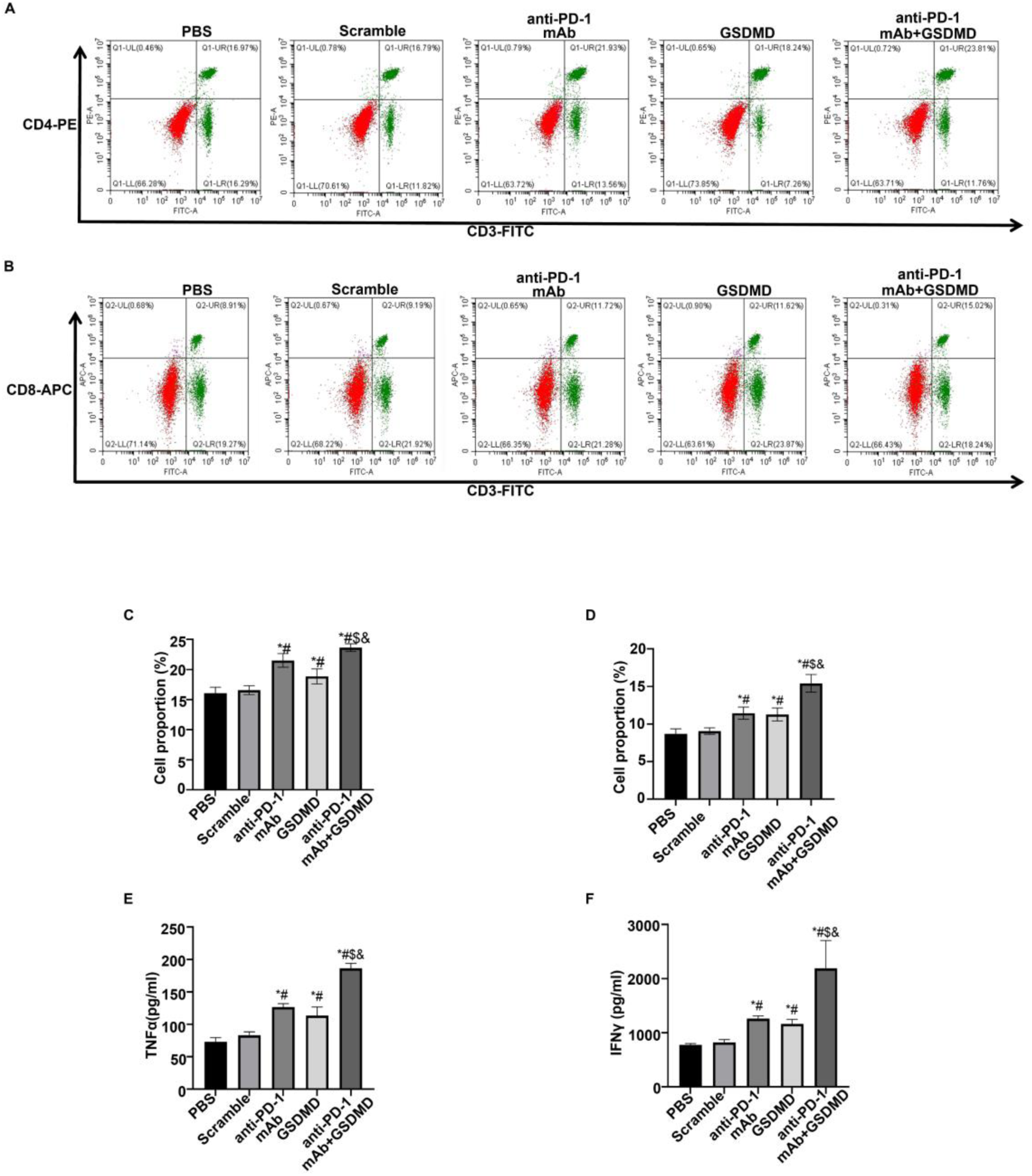
Combined therapy with GSDMD-expressing attenuated Salmonella and PD-1 blockade modulates peripheral T cell proportions and serum cytokine concentrations in tumor-bearing mice (A, C) Flow cytometric measurement and statistical analysis of CD4⁺ T cell percentages in peripheral blood. (B, D) Flow cytometry and quantitative statistics of peripheral CD8⁺ T cell proportions. (E) ELISA detection of serum TNF-α concentrations after combinatorial treatment. (F) ELISA analysis of serum IFN-γ levels across all experimental groups. \**P* < 0.05 versus PBS group; ^#^*P* < 0.05 versus Scramble group; ^$^*P* < 0.05 versus anti-PD-1 monotherapy group; ^&^*P* < 0.05 versus GSDMD monotherapy group.

CD8⁺ T cell proportions exhibited similar variation trends. To be specific, mice receiving combined therapy harbored the highest peripheral CD8⁺ T cell proportion among all experimental groups (Figure 7B, 7D). As two pivotal pro-inflammatory cytokines, TNF-α and IFN-γ co-modulate immune activation within the tumor microenvironment. An ELISA assay was further conducted to detect serum TNF-α and IFN-γ concentrations. The results showed that monotherapy significantly increased serum TNF-α and IFN-γ contents compared with PBS and Scramble groups, and the highest levels of both cytokines were observed in the combination group (Figure 7E–7F).

## 4. Discussion

Melanoma is an aggressive cutaneous malignancy that causes thousands of deaths annually. Despite tremendous advances in diagnostic screening and clinical management, the global incidence and mortality burden of melanoma continue to rise year by year [24]. Currently, exploring efficient therapeutic strategies for melanoma remains a core research hotspot and unmet clinical demand in oncology. Tumor immunotherapy, which suppresses tumor progression by modulating host immune homeostasis and activating immune cell functions, has emerged as a promising anti-cancer modality. In particular, mature immunotherapies represented by immune checkpoint inhibitors have been widely applied in clinical treatment [25,26]. Nevertheless, monotherapy targeting a single immune checkpoint suffers inherent therapeutic limitations, which are mainly attributed to tumor heterogeneity, differential target expression and therapy-induced drug resistance. To overcome such bottlenecks, novel combination regimens based on multi-target intervention have been continuously developed and translated into clinical practice, aiming to halt or delay tumor progression and optimize clinical therapeutic outcomes [27–29].

Herein, our in vivo experiments demonstrated that combinatorial administration of GSDMD-expressing attenuated Salmonella and anti-PD-1 antibody exerted superior anti-melanoma efficacy compared with monotherapy. This co-treatment substantially retarded melanoma growth, remodeled the immunosuppressive tumor microenvironment, and potentiated systemic anti-tumor immune responses in tumor-bearing mice.

Mechanistically, PD-1 binds to its ligand PD-L1 expressed on tumor cells, thereby triggering target cell apoptosis, restraining cytokine secretion and proliferation of PD-1-positive immune cells, and ultimately impairing the immune system’s capacity to recognize and eliminate malignant cells. Accordingly, the PD-1/PD-L1 signaling axis acts as a central pathway modulating tumor immune evasion and serves as a vital therapeutic target for cancer treatment [30,31]. Accumulating studies have validated that PD-1 blockade can reverse T cell exhaustion, facilitate the intratumoral infiltration of functional T lymphocytes, remodel local tumor immune microenvironment, and eventually amplify systematic anti-tumor immunity [32,33]. Consistent with these findings, our in vivo results verified that anti-PD-1 treatment not only elevated splenic T lymphocyte proportions to optimize systemic immune homeostasis, but also boosted the infiltration of CD4⁺ helper T cells and CD8⁺ cytotoxic T cells into melanoma lesions, thereby reshaping tumor immune landscape and triggering robust anti-tumor immune activation.

Additionally, pyroptosis refers to a lytic type of programmed cell death, which relies on caspase-mediated cleavage of gasdermin (GSDM) proteins to form plasma membrane pores [34]. As the most extensively characterized member of the GSDM family, GSDMD dominates pyroptosis execution. In the canonical pyroptosis pathway, inflammasome activation triggers caspase-1-mediated GSDMD cleavage, which further provokes robust inflammatory responses and ultimately initiates pyroptosis [35]. Accumulating evidence has demonstrated that targeting GSDMD represents a promising anti-tumor therapeutic strategy. It has been validated that 6,7-dichloro-2-methanesulfonyl-3-N-tert-butylaminoquinoxaline (DMB) acts as a direct and specific GSDMD agonist. DMB activates GSDMD pore-forming activity to trigger pyroptosis and suppress tumor growth, while causing negligible damage to GSDMD-expressing immune cells, thereby supporting the clinical feasibility of low-toxicity pyroptosis-targeted therapy [36].

￼

Existing melanoma-relevant studies have also verified that upregulated GSDMD expression restrains the proliferation, migration and invasion capacities of melanoma cells, indicating that GSDMD serves as a prospective therapeutic target for melanoma treatment [37]. Notably, mounting studies have demonstrated that GSDMD-dependent pyroptosis is capable of triggering protective anti-tumor immune responses. Inflammatory factors released during GSDMD-mediated pyroptosis facilitate intratumoral CD8⁺ T cell infiltration, reverse tumor immune evasion, and thereby block gastric cancer progression [38]. Another study reported that inactivation of mixed-lineage leukemia 4 (MLL4)-dependent enhancers induces GSDMD-related pyroptosis, so as to amplify anti-tumor immunity [39]. Moreover, stimulator of interferon genes (STING) targeting elicits GSDMD-dependent pyroptosis and potentiates immunotherapeutic efficacy against renal cell carcinoma [40].

Consistent with the above published findings, our in vivo outcomes confirmed that GSDMD-delivering attenuated Salmonella efficiently boosted intratumoral GSDMD expression and activated host anti-tumor immunity. Specifically, GSDMD engineering elevated splenic T lymphocyte proportions and enhanced the intratumoral infiltration of CD4⁺ helper T cells and CD8⁺ cytotoxic T cells. Most importantly, compared with monotherapy, combined treatment with GSDMD-expressing Salmonella and anti-PD-1 antibody exerted superior immunomodulatory effects. The combinatorial regimen achieved the most significant improvements in both splenic T lymphocyte abundance and tumoral T-cell infiltration among all experimental groups.

Unquestionably, as the core effector protein that assembles functional pores on the plasma membrane, GSDMD serves as an indispensable executor of pyroptosis. Accumulating data have also revealed that upregulated GSDMD expression elevates cellular apoptotic rates and participates in the coordinated regulation of multiple cell death programs [41,42]. Similarly, PD-1 blockade reverses T cell exhaustion to drive robust T cell activation. Upon activation, T lymphocytes secrete abundant pro-inflammatory cytokines including IL-2, tumor necrosis factor-α (TNF-α) and interferon-γ (IFN-γ). These cytokines cooperatively target malignant cells and facilitate programmed tumor cell death by modulating pro-apoptotic signaling cascades [43].

Consistent with these published mechanistic findings, our in vivo data demonstrated that monotherapy with either anti-PD-1 antibody or GSDMD-expressing attenuated Salmonella effectively augmented intratumoral apoptotic activity, conferring independent pro-apoptotic anti-tumor effects. Notably, combined treatment generated a prominent synergistic pro-apoptotic response; the apoptotic index in the co-treatment group was markedly higher than that of either monotherapy cohort, confirming a synergistic mechanism linking PD-1 immune checkpoint inhibition and GSDMD-mediated pyroptosis in triggering melanoma cell apoptosis.

Collectively, the present study validated that combinatorial intervention with GSDMD-delivering attenuated Salmonella and PD-1 blockade elicits superior anti-melanoma efficacy via facilitating melanoma cell apoptosis and potentiating systemic anti-tumor immune responses in tumor-bearing mice. Our findings provide preclinical evidence supporting the development of combinatorial regimens for melanoma management.

Nevertheless, conflicting reports regarding the dual role of GSDMD in tumor progression have been documented. For instance, one study illustrated that gut microbiota-driven NLRP3-dependent GSDMD activation accelerates colorectal tumorigenesis. Genetic ablation of Gsdmd markedly suppresses colorectal tumor formation in mice, indicating a pro-tumorigenic function of GSDMD in intestinal malignancies [44]. Accordingly, the precise molecular mechanisms underlying the anti-melanoma activity of GSDMD delivered by attenuated Salmonella remain to be fully elucidated. Furthermore, critical therapeutic parameters for this combinatorial regimen—including the optimal treatment timing window, matched dosage ratio and administration frequency—require systematic optimization and validation in follow-up investigations. These unresolved questions constitute the primary focus of our future research, which will lay a more solid experimental foundation to advance the clinical translation of this synergistic therapeutic strategy.

## Supporting information

One of the Raw western blot results for Figure 3

One of the Raw western blot results for Figure 3

One of the Raw western blot results for Figure 3

One of the Raw western blot results for Figure 3

One of the Raw western blot results for Figure 6G

One of the Raw western blot results for Figure 6G

## Acknowledgments

This study was financially supported by the Xinxiang University Doctoral Start-up Grant (grant nos. 1366020279) awarded to Xiaolong Jia, the Henan Provincial Young and Middle-aged Health Science and Technology Innovation Talents Training Project (grant nos. LJRC2023013) awarded to Lei Wang, the Technology Research Project of Henan Province (grant nos. 242102310136) awarded to Lei Wang.

## References

[1] Druskovich C, Kelley J, Aubrey J, Palladino L, Wright G P. A Review of Melanoma Subtypes: Genetic and Treatment Considerations[J]. Journal of surgical oncology, 2025,131:356–364.

[2] Rumgay H, Nethan S T, Shah R, Vignat J, Ayo-Yusuf O, Chaturvedi P, et al. Global burden of oral cancer in 2022 attributable to smokeless tobacco and areca nut consumption: a population attributable fraction analysis[J]. The Lancet. Oncology, 2024,25:1413–1423.

[3] Kraft T, Grützmann K, Meinhardt M, Meier F, Westphal D, Seifert M. Personalized identification and characterization of genome-wide gene expression differences between patient-matched intracranial and extracranial melanoma metastasis pairs[J]. Acta neuropathologica communications, 2024,12:67.

[4] Xiong L, Cheng J. Rewiring lipid metabolism to enhance immunotherapy efficacy in melanoma: a frontier in cancer treatment[J]. Frontiers in oncology, 2025,15:1519592.

[5] Arafat Hossain M. A comprehensive review of immune checkpoint inhibitors for cancer treatment[J]. International immunopharmacology, 2024,143:113365.

[6] Carlino M S, Larkin J, Long G V. Immune checkpoint inhibitors in melanoma[J]. Lancet (London, England), 2021,398:1002–1014.

[7] Mallardo D, Basile D, Vitale M G. Advances in Melanoma and Skin Cancers[J]. International journal of molecular sciences, 2025,26.

[8] Jiang Y, Chen M, Nie H, Yuan Y. PD-1 and PD-L1 in cancer immunotherapy: clinical implications and future considerations[J]. Human vaccines & immunotherapeutics, 2019,15:1111–1122.

[9] Liu X, Zhao A, Xiao S, Li H, Li M, Guo W, et al. PD-1: A critical player and target for immune normalization[J]. Immunology, 2024,172:181–197.

[10] Dong Y, Sun Q, Zhang X. PD-1 and its ligands are important immune checkpoints in cancer[J]. Oncotarget, 2017,8:2171–2186.

[11] Jiang X, Wang J, Deng X, Xiong F, Ge J, Xiang B, et al. Role of the tumor microenvironment in PD-L1/PD-1-mediated tumor immune escape[J]. Molecular cancer, 2019,18:10.

[12] Keir M E, Butte M J, Freeman G J, Sharpe A H. PD-1 and its ligands in tolerance and immunity[J]. Annual review of immunology, 2008,26:677–704.

[13] Homet Moreno B, Parisi G, Robert L, Ribas A. Anti-PD-1 therapy in melanoma[J]. Seminars in oncology, 2015,42:466–473.

[14] Habibi M A, Mirjani M S, Ahmadvand M H, Delbari P, Eftekhar M S, Ghazizadeh Y, et al. Anti-PD-1/PD-L1 inhibitor therapy for melanoma brain metastases: a systematic review and meta-analysis[J]. Neurosurgical review, 2024,47:434.

[15] Tsai K K, Zarzoso I, Daud A I. PD-1 and PD-L1 antibodies for melanoma[J]. Human vaccines & immunotherapeutics, 2014,10:3111–3116.

[16] Du F, Yang L H, Liu J, Wang J, Fan L, Duangmano S, et al. The role of mitochondria in the resistance of melanoma to PD-1 inhibitors[J]. Journal of translational medicine, 2023,21:345.

[17] Dong W W, Liu T, He L X, He W T. Targeting pyroptosis in inflammatory bowel disease: A potentially effective therapeutic approach[J]. World journal of gastroenterology, 2025,31:111358.

[18] Johnson D E, Cui Z. Triggering Pyroptosis in Cancer[J]. Biomolecules, 2025,15.

[19] Wu D, Zhang Y, Guo Y, Dong Z. Pyroptosis as a molecular bridge: linking gasdermin activation to enhanced anti-tumor immunity in immunotherapy[J]. Biochemical pharmacology, 2026,244:117587.

[20] Chen G, Zhang Z, Chong W, Qiu G, Li Z, Yang S, et al. The molecular mechanisms of pyroptosis and its implications in tumor immunotherapy[J]. Molecular cancer, 2026.

[21] Ortega-Ribera M, Zhuang Y, Brezani V, Joshi R S, Zsengeller Z, Nagesh P T, et al. Gasdermin D deletion prevents liver injury and exacerbates extrahepatic damage in a murine model of alcohol-induced ACLF[J]. eGastroenterology, 2025,3:e100151.

[22] Yuan R, Zhao W, Wang Q Q, He J, Han S, Gao H, et al. Cucurbitacin B inhibits non-small cell lung cancer in vivo and in vitro by triggering TLR4/NLRP3/GSDMD-dependent pyroptosis[J]. Pharmacological research, 2021,170:105748.

[23] Chen J, Xu M, Wu F, Wu N, Li J, Xie Y, et al. CRKL silencing inhibits melanoma growth and enhances its chemotherapy sensitivity through the PI3K/AKT and NLRP3/GSDMD pathways[J]. Biochemical pharmacology, 2025,235:116840.

[24] Shah M, Schur N, Rosenberg A, DeBusk L, Burshtein J, Zakria D, et al. Trends in Melanoma Incidence and Mortality[J]. Dermatologic clinics, 2025,43:373–379.

[25] Dillman R O. Cancer immunotherapy[J]. Cancer biotherapy & radiopharmaceuticals, 2011,26:1–64.

[26] Zhang M, Liu C, Tu J, Tang M, Ashrafizadeh M, Nabavi N, et al. Advances in cancer immunotherapy: historical perspectives, current developments, and future directions[J]. Molecular cancer, 2025,24:136.

[27] Damia G, Broggini M. Editorial: Innovative drug combinations for enhanced solid tumor treatment efficacy[J]. Frontiers in oncology, 2025,15:1763808.

[28] Doostmohammadi A, Jooya H, Ghorbanian K, Gohari S, Dadashpour M. Potentials and future perspectives of multi-target drugs in cancer treatment: the next generation anti-cancer agents[J]. Cell communication and signaling : CCS, 2024,22:228.

[29] Damodaran C, Cho J Y, Güngör C. Therapeutic resistance and combination therapy for cancer: recent developments and future directions[J]. Scientific reports, 2025,15:26881.

[30] Xu P, Hong C, Liu L, Xiao L. PD-1/PD-L1 blockade therapy in hepatocellular carcinoma: Current status and potential biomarkers[J]. Biochimica et biophysica acta. Reviews on cancer, 2025,1880:189334.

[31] Syamsu S A, Faruk M, Smaradania N, Sampepajung E, Pranoto A S, Irsandy F, et al. PD-1/PD-L1 pathway: Current research in breast cancer[J]. Breast disease, 2024,43:79–92.

[32] Kikuchi H, Matsui A, Morita S, Amoozgar Z, Inoue K, Ruan Z, et al. Increased CD8+ T-cell Infiltration and Efficacy for Multikinase Inhibitors After PD-1 Blockade in Hepatocellular Carcinoma[J]. Journal of the National Cancer Institute, 2022,114:1301–1305.

[33] Lee A H, Sun L, Mochizuki A Y, Reynoso J G, Orpilla J, Chow F, et al. Neoadjuvant PD-1 blockade induces T cell and cDC1 activation but fails to overcome the immunosuppressive tumor associated macrophages in recurrent glioblastoma[J]. Nature communications, 2021,12:6938.

[34] Broz P, Pelegrín P, Shao F. The gasdermins, a protein family executing cell death and inflammation[J]. Nature reviews. Immunology, 2020,20:143–157.

[35] Zhang N, Zhang J, Yang Y, Shan H, Hou S, Fang H, et al. A palmitoylation-depalmitoylation relay spatiotemporally controls GSDMD activation in pyroptosis[J]. Nature cell biology, 2024,26:757–769.

[36] Fontana P, Du G, Zhang Y, Zhang H, Vora S M, Hu J J, et al. Small-molecule GSDMD agonism in tumors stimulates antitumor immunity without toxicity[J]. Cell, 2024,187:6165–6181.e6122.

[37] Zhao S, Zhu Y, Liu H, He X, Xie J. System analysis based on the pyroptosis-related genes identifes GSDMD as a novel therapy target for skin cutaneous melanoma[J]. Journal of translational medicine, 2023,21:801.

[38] Kang M, Du W, Ding L, Wu M, Pei D. HIC1 suppresses Tumor Progression and Enhances CD8(+) T Cells Infiltration Through Promoting GSDMD-induced Pyroptosis in Gastric Cancer[J]. Advanced science (Weinheim, Baden-Wurttemberg, Germany), 2025,12:e2412083.

[39] Ning H, Huang S, Lei Y, Zhi R, Yan H, Jin J, et al. Enhancer decommissioning by MLL4 ablation elicits dsRNA-interferon signaling and GSDMD-mediated pyroptosis to potentiate anti-tumor immunity[J]. Nature communications, 2022,13:6578.

[40] Wu S, Wang B, Li H, Wang H, Du S, Huang X, et al. Targeting STING elicits GSDMD-dependent pyroptosis and boosts anti-tumor immunity in renal cell carcinoma[J]. Oncogene, 2026,45:620–635.

[41] Shah S S, Manning J A, Lim Y, Sinha D, Murthy A M V, Ganesan R, et al. NEDD4L-mediated Gasdermin D and E ubiquitination regulates cell death and tissue injury[J]. Cell death and differentiation, 2025.

[42] Zhu C L, Xie J, Liu Q, Wang Y, Li H R, Yu C M, et al. PD-L1 promotes GSDMD-mediated NET release by maintaining the transcriptional activity of Stat3 in sepsis-associated encephalopathy[J]. International journal of biological sciences, 2023,19:1413-1429.

[43] Moseman J E, Rastogi I, Jeon D, McNeel D G. PD-1 blockade employed at the time CD8+ T cells are activated enhances their antitumor efficacy[J]. Journal for immunotherapy of cancer, 2025,13.

[44] Chen J, Singh N, Ye X, Theune E V, Wang K. Gut microbiota-mediated activation of GSDMD ignites colorectal tumorigenesis[J]. Cancer gene therapy, 2024,31:1007–1017.

