## Supplementary figures and images for "Attenuated Salmonella-Mediated Delivery of GSDMD Potentiates PD-1 Blockade Therapy against Melanoma"

### One of the Raw western blot results for Figure 3

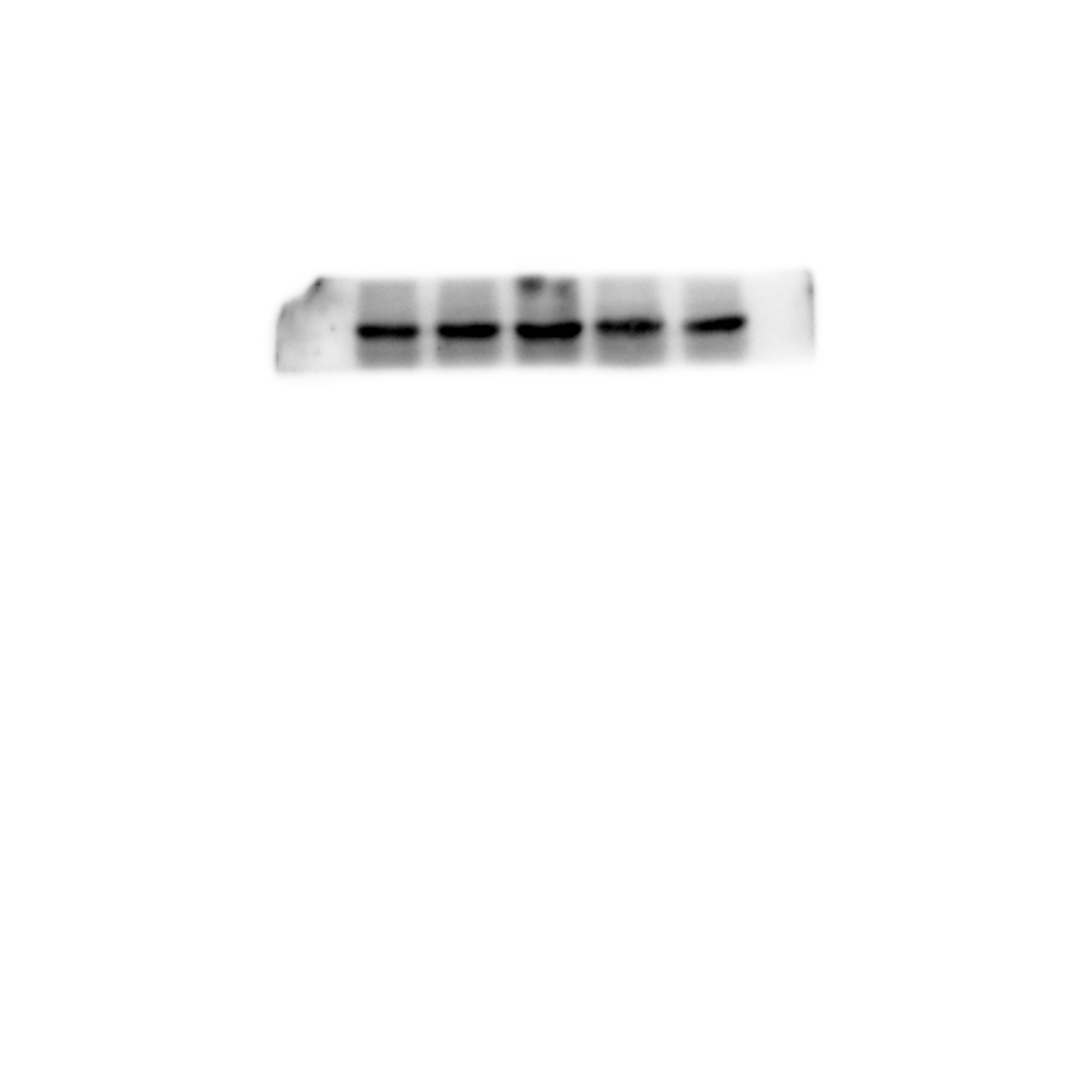

### One of the Raw western blot results for Figure 3

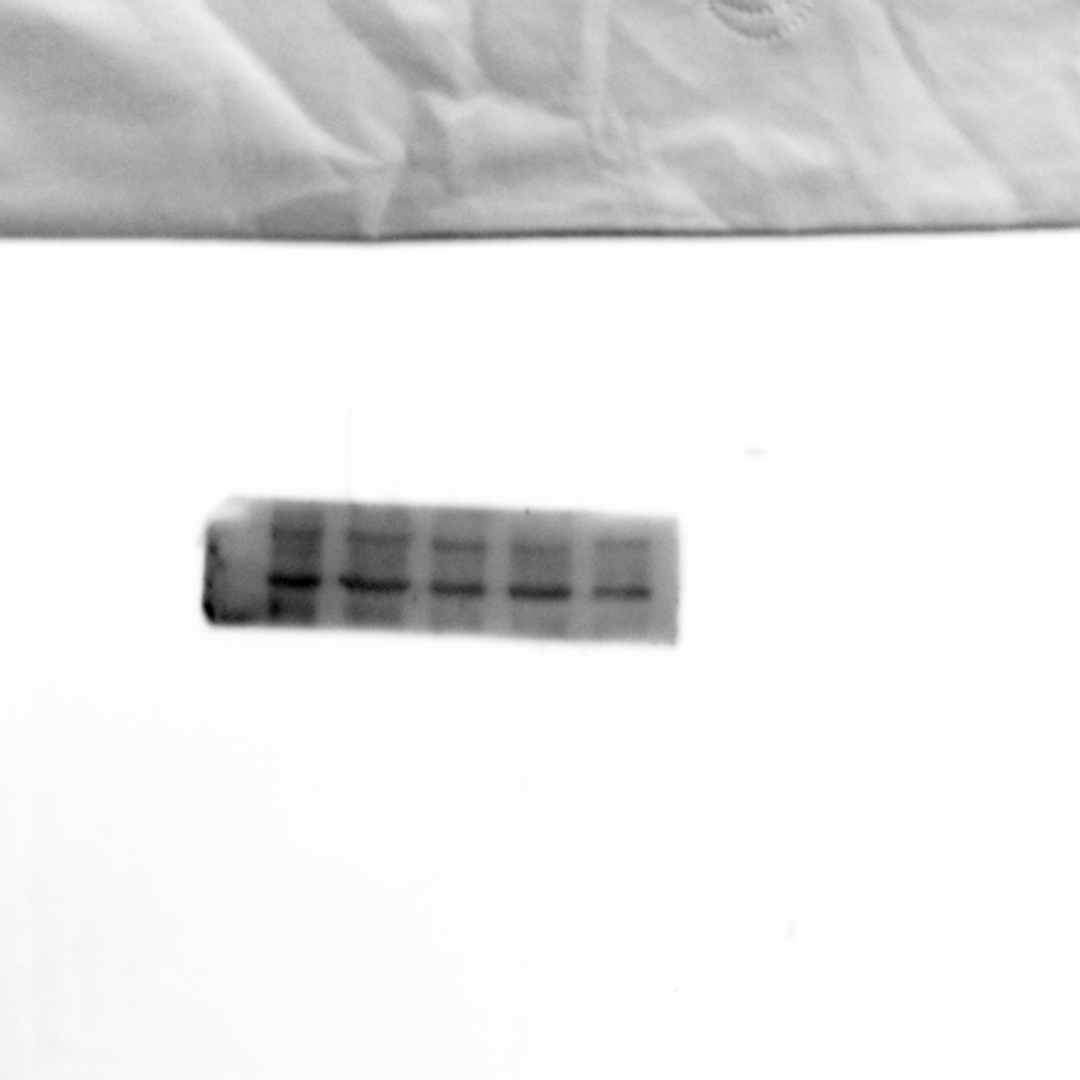

### One of the Raw western blot results for Figure 3

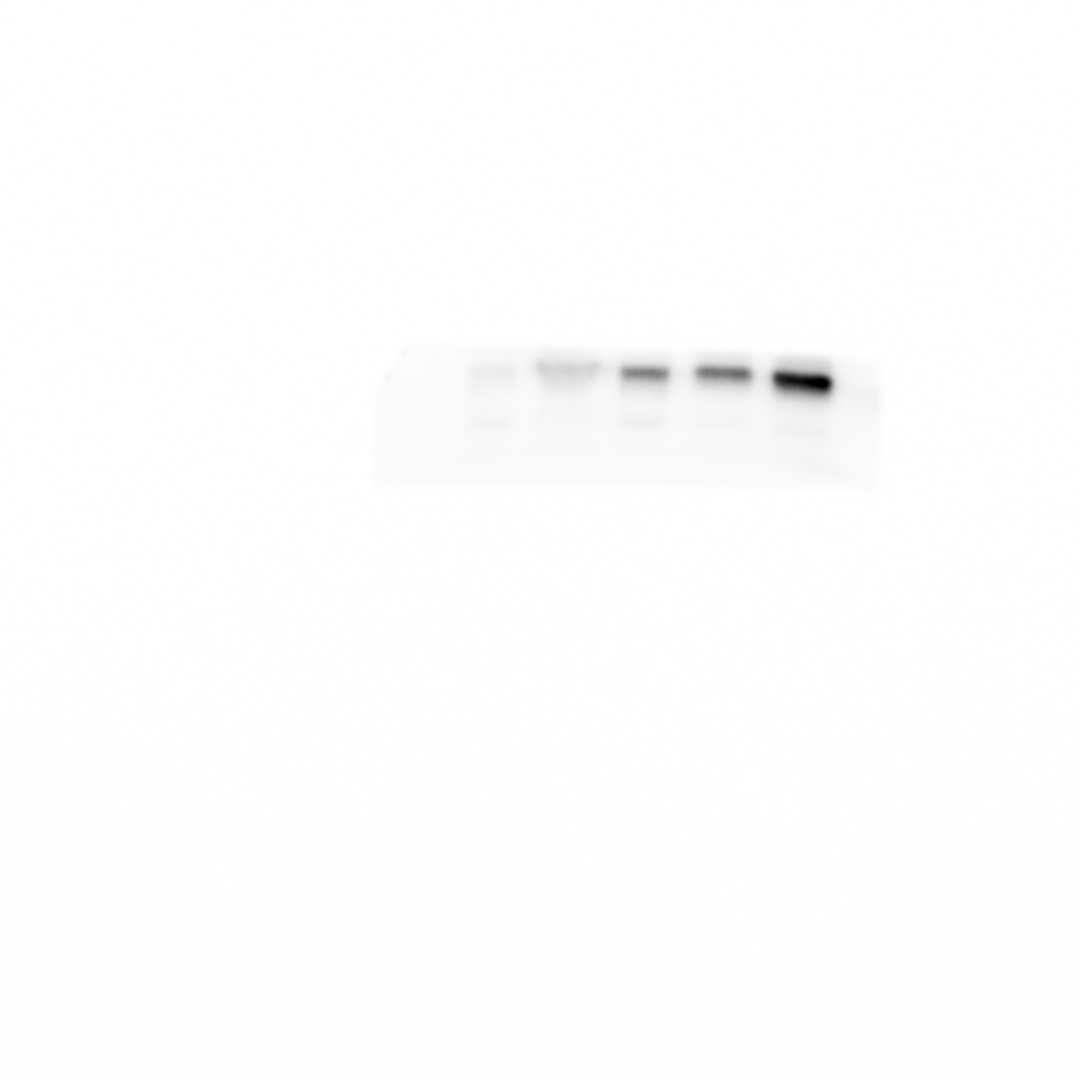

### One of the Raw western blot results for Figure 3

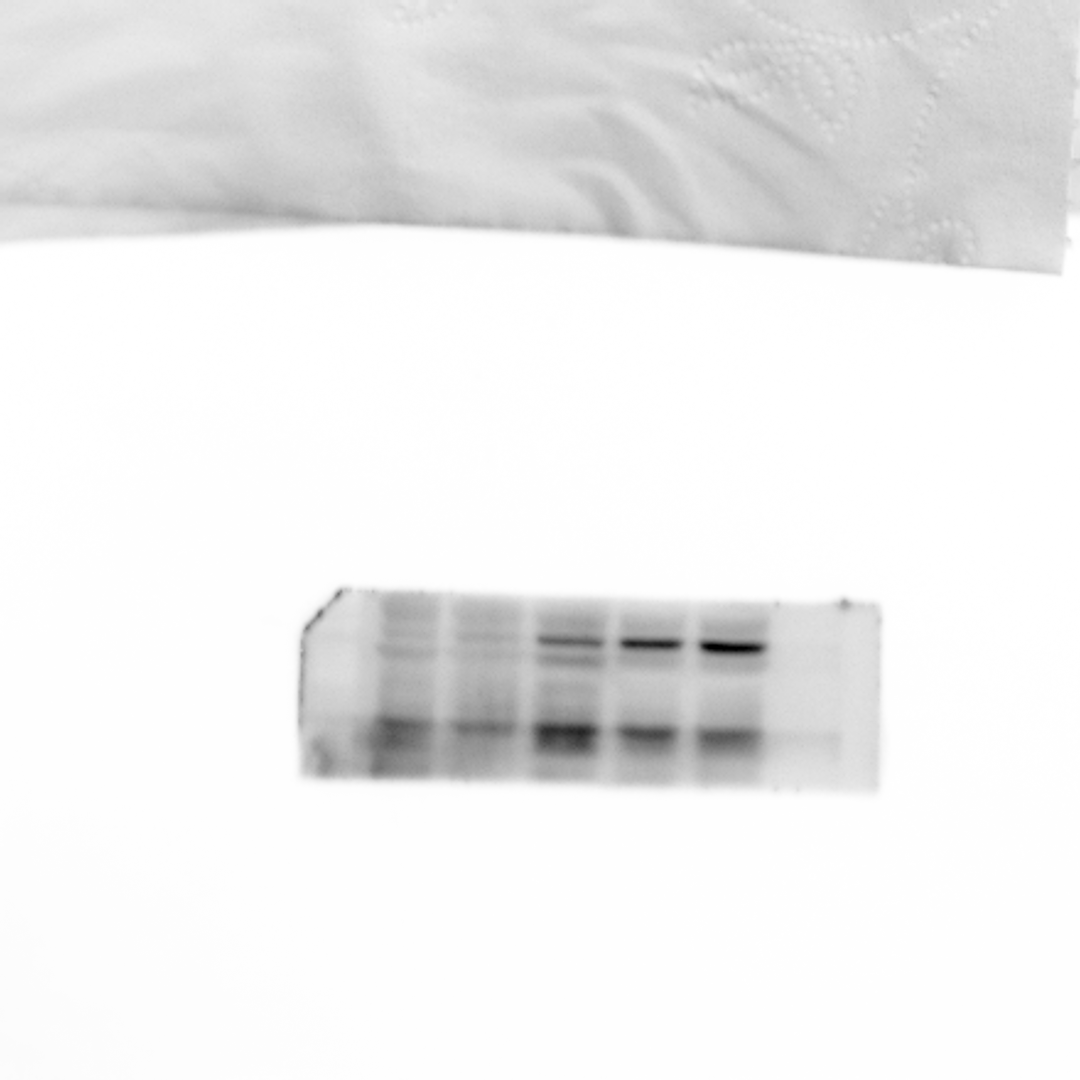

### One of the Raw western blot results for Figure 6G

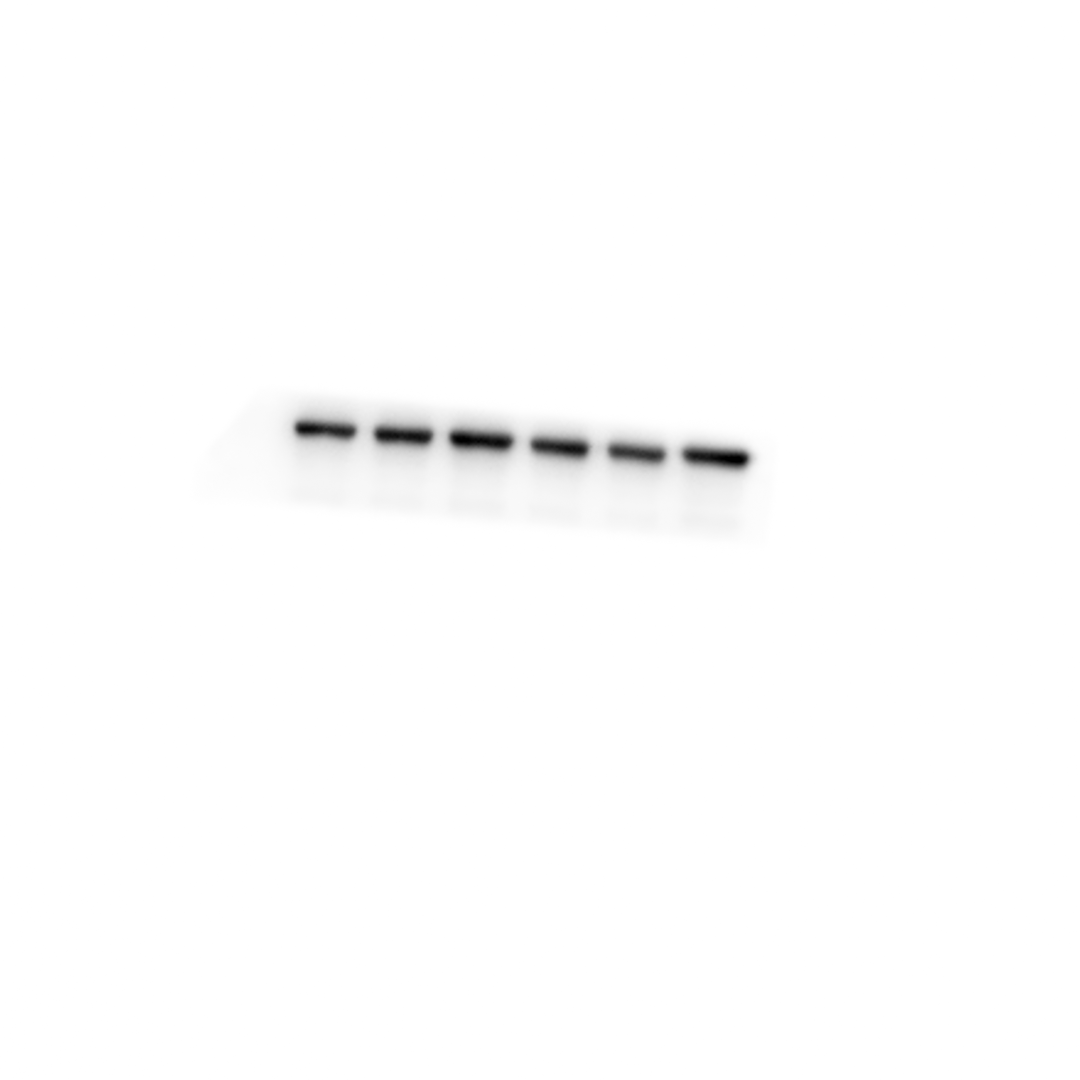

### One of the Raw western blot results for Figure 6G

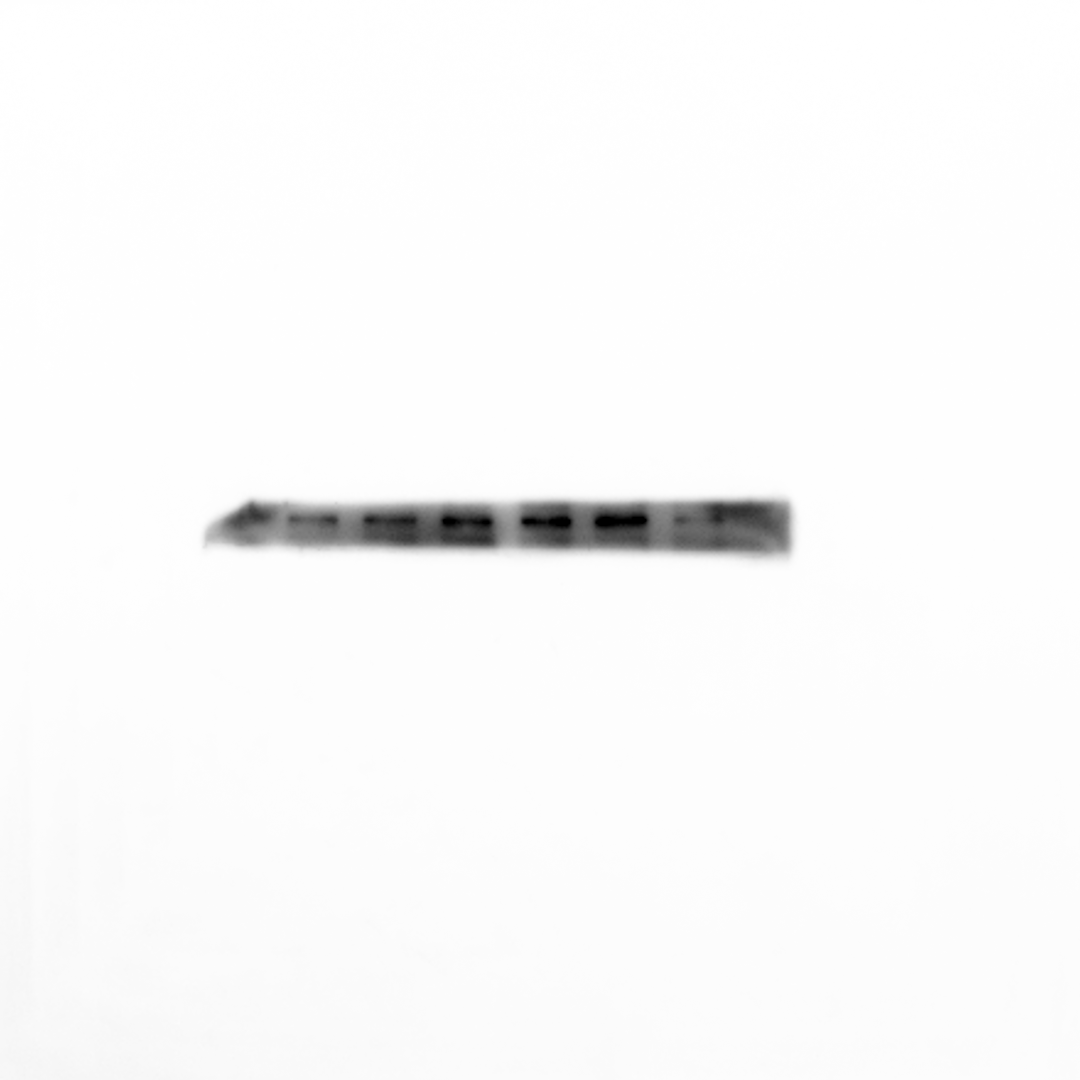
